# Globally unified analysis of riverine eDNA reveals common associations of fish biodiversity with drainage characteristics

**DOI:** 10.1101/2025.09.24.676958

**Authors:** Yan Zhang, Heng Zhang, Hiroshi Akashi, Camille P. Albouy, Kara J. Andres, José Barquín, Jeanine Brantschen, Richard E. Connon, Joseph M. Craine, Deirdre B. Gleeson, Alejandra Goldenberg-Vilar, Alexia M. González-Ferreras, Chelsea Hatzenbuhler, Kamil Hupało, Josephine Hyde, Wataru Iwasaki, Mark D. Johnson, Aron D. Katz, Vyacheslav V. Kuzovlev, Courtney E. Larson, Laurène A. Lecaudey, Florian Leese, Matthieu Leray, Feilong Li, Till-Hendrik Macher, Quentin Mauvisseau, María Morán-Luis, Georgia Nester, Helio Quintero, Tsilavina Ravelomanana, Merin Reji Chacko, Mattia Saccò, Naiara Sales, Tamara Schenekar, Martin Schletterer, Saskia Schmidt, Nicholas O. Schulte, Robin Schütz, Jinelle H. Sperry, Emma R. Stevens, Sarah A. Stinson, Steven Weiss, Fei Xia, Hui Zhang, Song Zhang, Wenjun Zhong, Shuo Zong, Loïc Pellissier, Xiaowei Zhang, Florian Altermatt

## Abstract

Freshwater biodiversity is declining at a pace that outstrips the capacity of existing monitoring approaches both in temporal and spatial dimensions, highlighting the urgent need for rapid and scalable assessment and attribution of biodiversity states and changes. Here, we present one of the first global assessments and unified analyses of riverine fish biodiversity using environmental DNA (eDNA) collected from 1818 sites across 113 river systems. We quantified species richness, functional redundancy, phylogenetic diversity, and genetic sequence diversity, and related them to drainage characteristics. Our results showed that eDNA effectively captured global patterns of multi-faceted riverine fish biodiversity and disentangled the roles of climate and human activities in shaping biodiversity–area relationships. Catchments in warmer climates consistently enhanced biodiversity accumulation with area, while higher human activity intensity weakened this scaling. Species richness, functional, and genetic sequence diversity exhibited stronger negative responses to human activities in larger catchments. In contrast, phylogenetic diversity showed the strongest negative effects in smaller catchments with these impacts diminishing as catchment area increased, highlighting the facet-dependent nature of biodiversity responses to environmental gradients. Our findings demonstrate the power of eDNA-based datasets for harmonized, multi-faceted biodiversity assessments, offering a scalable approach for detecting and attributing biodiversity change and informing conservation strategies under accelerating global change.

## Introduction

Global biodiversity is declining at an unprecedented rate, while efforts in biodiversity conservation and ecosystem restoration are lagging behind^1–3^. Next to declines in richness and population sizes, the erosion of within-population genetic diversity is becoming increasingly apparent, with losses occurring worldwide under the influence of human activities^4,5^. Freshwater ecosystems cover less than 1% of Earth’s surface but sustain nearly 10% of known species, underscoring their disproportionate importance to global biodiversity^6^. Yet, freshwater ecosystems are also among the most threatened biomes, with population declines of up to 85% for freshwater species^4,7,8^. These declines are particularly strong in freshwater fishes, which are both highly vulnerable^9,10^, yet also particularly important with respect to ecosystem services. The declines are jointly driven by multiple pressures including climate change, land-use transformation, overexploitation, and biological invasions, which have profoundly reshaped species distributions, community structures, and ecosystem functions^11^.

For effective conservation and mitigation of these challenges, there is not only a need to understand the drivers themselves, but also to understand their effects in the context of universal physical structures and properties of river networks^12^, which shape the distribution and persistence of riverine biodiversity. River networks exhibit hierarchical branching structures that constrain the movement of organisms, resources, and energy^12–14^. This dendritic configuration embeds river systems within the landscape, where upstream flows transport biotic and abiotic signals that shape downstream ecosystems^15,16^. Biodiversity is generally expected to increase from headwaters to mainstems due to stronger connectivity and cumulative species inflow, following a typical biodiversity–area relationship^17^. However, these patterns are not static. Climatic variation and human activities interact with hydrological processes, altering biodiversity patterns by reshaping community structure and diversity patterns along the river continuum^18,19^. Such spatial heterogeneity in responses to human activities may partly explain why variation in biodiversity patterns has been observed across individual catchments worldwide^20,21^. Variable patterns may also partially reflect the complex interplay between river network structure, climatic gradients, and human activities. Yet, past work on riverine diversity is often based on very different monitoring and assessment approaches, and the universality of local to global riverine diversity patterns is subject for debate^22,23^.

Traditionally, biodiversity has been looked at from a species richness perspective. However, variation in functional, phylogenetic, and genetic diversity are further dimensions of diversity capturing complementary aspects of ecosystem structure, function and stability^10,24^. Functional diversity reflects ecological niche differentiation and functional robustness, with functional redundancy enhancing ecosystem resilience to species loss^25^. Phylogenetic diversity describes the evolutionary distinctiveness of the community ^26^. Finally, genetic diversity directly captures the adaptive potential of populations, which in turn reduces population fluctuations and extinction risk, ultimately enhancing the long-term functional stability of ecosystems^5,27^. Despite the growing emphasis on monitoring biodiversity across multiple facets, as for example advocated by the Global Biodiversity Framework (GBF)^28^, comprehensive assessments of the spatial patterns and drivers of multi-faceted biodiversity in riverine ecosystems are largely lacking.

In recent years, environmental DNA (eDNA) has emerged as a transformative tool for aquatic biodiversity monitoring and has been widely applied across ecosystems^29^. Environmental DNA approaches capture taxonomic diversity^30,31^, enable estimation of functional diversity and link to species’ trait data^32,33^, calculation of phylogenetic diversity and, to some extent, assessment of genetic diversity based on sequence information^34,35^. As such, it has become the method of choice for rapid and quantitative assessment of multiple facets of biodiversity at larger spatial scales^36,37^, possibly enabling a scalable and spatially representative biodiversity monitoring and observation networks^38^. While the effectiveness of eDNA has been well validated at regional scales, most existing applications remain focused on specific regions or individual facets of biodiversity^16,39,40^. Recently, the use of aquatic eDNA to address global biodiversity targets has been postulated^41^, yet requires a demonstration of consistent and robust assessment of multidimensional biodiversity patterns across catchment basins, climate zones, and gradients of human activities.

Here, we analyze multi-faceted riverine fish biodiversity patterns, covering species, functional, phylogenetic, and genetic diversity, in a unifying framework and link these diversity patterns to natural and human-related drivers. We build on the first globally unified and harmonized fish eDNA dataset^42^, integrating both original individually-reported and unified community data from 113 river catchments across six continents and encompasses 1818 individual sampling sites, spanning tropical to temperate zones and address three key questions: (1) to what extent does eDNA capture globally relevant, macroecological patterns of riverine fish biodiversity (covering multiple facets including taxonomic, functional, phylogenetic, and genetic sequence diversity), and how consistent are these patterns when compared to conventional biodiversity data? (2) how do drainage characteristics, particularly climatic and human activities in combination with catchment area, modulate biodiversity–area relationships across river systems? (3) how sensitive is fish biodiversity to human activities across the river network, and are there generalizable differences between upstream and downstream locations or across drainage sizes? By combining biodiversity metrics derived from eDNA with drainage characteristics, our study provides one of the first global-scale tests of eDNA-based riverine biodiversity assessments, demonstrating how fish biodiversity responds to variation in drainage properties and human activities.

## Results

### Validation of eDNA-derived biodiversity

We characterized riverine fish diversity across four complementary biodiversity facets—species richness, functional redundancy, mean nearest taxon distance (MNTD), and genetic sequence diversity—based on a unified and standardized analysis of raw eDNA sequencing data (Figure 1). For comparison, we also calculated four corresponding metrics: species richness and functional redundancy from original individually-reported eDNA datasets, phylogenetic diversity represented as tree-based MNTD from a global fish phylogeny, and barcode-derived genetic sequence diversity derived from COI barcode sequences in NCBI.

**Figure 1.**
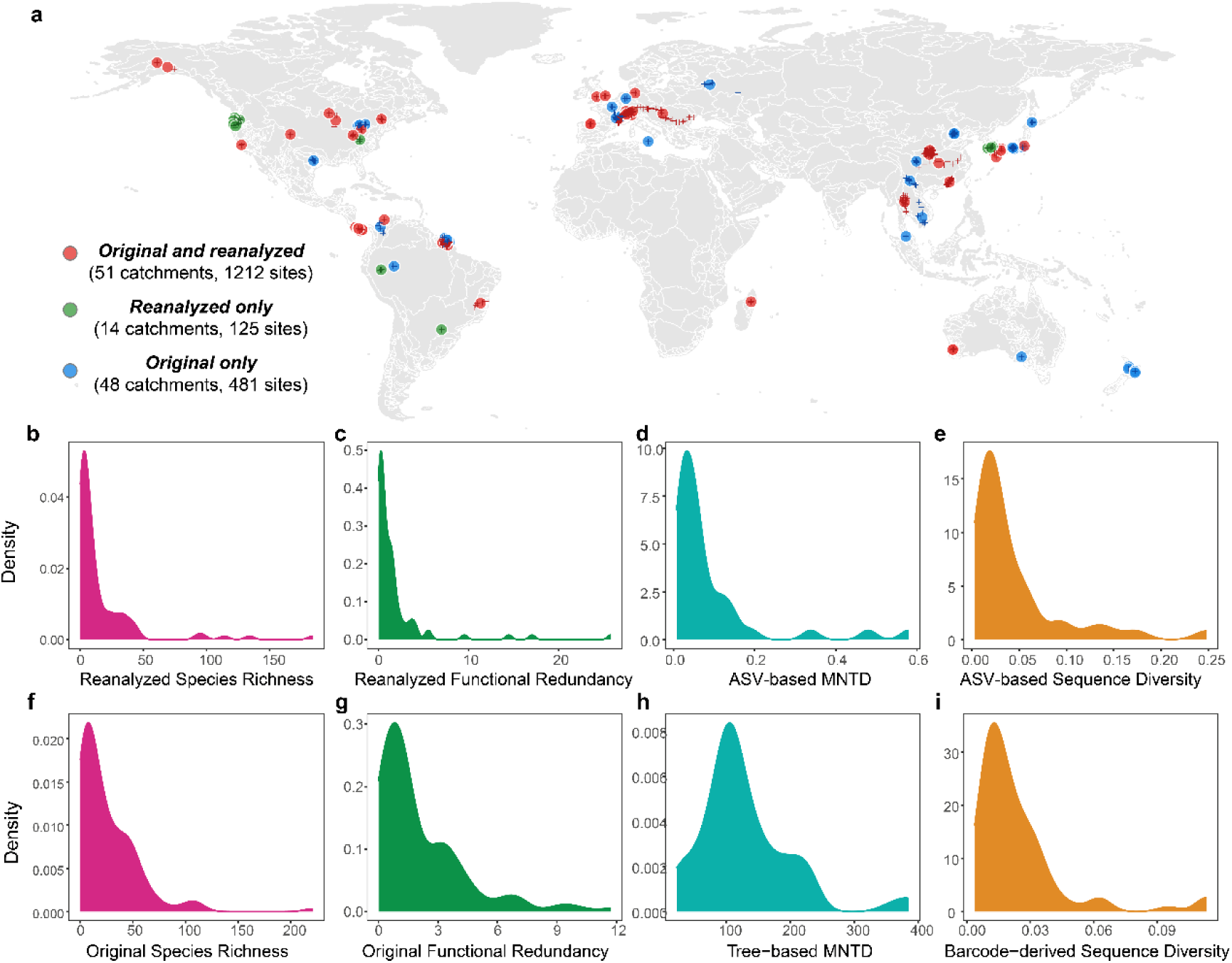
Global synthesis of fish diversity based on environmental DNA data. (a), Geographic distribution of 113 river catchments (circles), comprising a total of 1818 eDNA sampling sites included in this meta-analysis (sites are added as crosses). Fish community data were compiled from studies that either original individually-reported fish diversity metrics (blue), provided raw sequencing data for unified and standardized analysis (green), or both (red). The unified fish diversity facets derived from a standardized bioinformatic pipeline applied to raw eDNA sequencing data, including (b) species richness, (c) functional redundancy calculated from unified species-by-site tables using species-level trait information, (d) mean nearest taxon distance (MNTD) calculated from ASVs, and (e) genetic sequence diversity estimated from within-species nucleotide variation based on ASV sequences. Original individually-reported diversity metrics based on values from individual studies, including (f) species richness and, when available, (g) functional redundancy. (h) MNTD computed from unified species-by-site tables using a published global fish phylogeny to quantify phylogenetic relatedness among species. (i) Barcode-derived genetic sequence diversity calculated from unified species lists by retrieving COI sequences for each species from NCBI and computing within-species nucleotide diversity.

Across facets, correlation analyses among diversity facets revealed that species richness and functional redundancy were strongly correlated, whereas phylogenetic and genetic sequence diversity showed weaker associations with other facets (Supplementary Figure 1). Species richness and functional redundancy both exhibited clear tropical peaks and declined towards higher latitudes (Supplementary Figure 2: GAM R² = 0.18 and 0.16, respectively, *p* < 0.001). Phylogenetic diversity, in contrast, was higher at higher latitudes, particularly for tree-based MNTD, which showed a stronger positive relationship with latitude (R² = 0.14, *p* < 0.001). Genetic sequence diversity was generally highest in the tropics, especially when derived from COI barcodes (R² = 0.15, *p* < 0.001) with weaker signals for ASV-based estimates (R² = 0.08). Together, these results reveal marked spatial heterogeneity across taxonomic, functional, phylogenetic, and genetic dimensions of riverine fish diversity at the global scale.

To evaluate the reliability of eDNA-derived diversity metrics, we systematically compared them with conventional biodiversity records (Figure 2 and Supplementary Table 1). Unified species richness was highly correlated with historical basin-level species records, showing robust consistency in models that included either primer (GLMM, *p* < 0.001, marginal R² = 0.16, conditional R² = 0.70) or barcode (*p* < 0.001, marginal R² = 0.27, conditional R² = 0.51) as a random effect. Notably, these correlations were stronger than those for original individually reported species richness (Supplementary Figure 3, marginal R² = 0.08–0.10, conditional R² = 0.39–0.68). For phylogenetic diversity, alpha diversity estimated from ASV-based MNTD showed strong correlation with that from tree-based MNTD (*p* < 0.001, marginal R² = 0.22, conditional R² = 0.62; in models that included barcode as a random effect, *p* < 0.001, marginal R² = 0.19, conditional R² = 0.22), whereas gamma diversity showed notably weaker correlation (Supplementary Figure 3, *p* > 0.05, marginal R² = 0.04–0.05, conditional R² = 0.05–0.20). For genetic sequence diversity, gamma diversity of ASV-based within-species nucleotide diversity was marginally significantly correlated with barcode-derived estimates (Supplementary Figure 3, *p* = 0.094, marginal R² = 0.08, conditional R² = 0.08), while alpha diversity showed a weaker but highly significant correlation in models that included barcode as a random effect (*p* < 0.001, marginal R² = 0.02, conditional R² = 0.19). This pattern was consistent across genetic sequence diversity estimates derived from both 12S and 16S barcode sequences (Supplementary Figure 4), demonstrating the robustness of ASV-based genetic sequence diversity relative to traditional barcode-based approaches.

**Figure 2.**
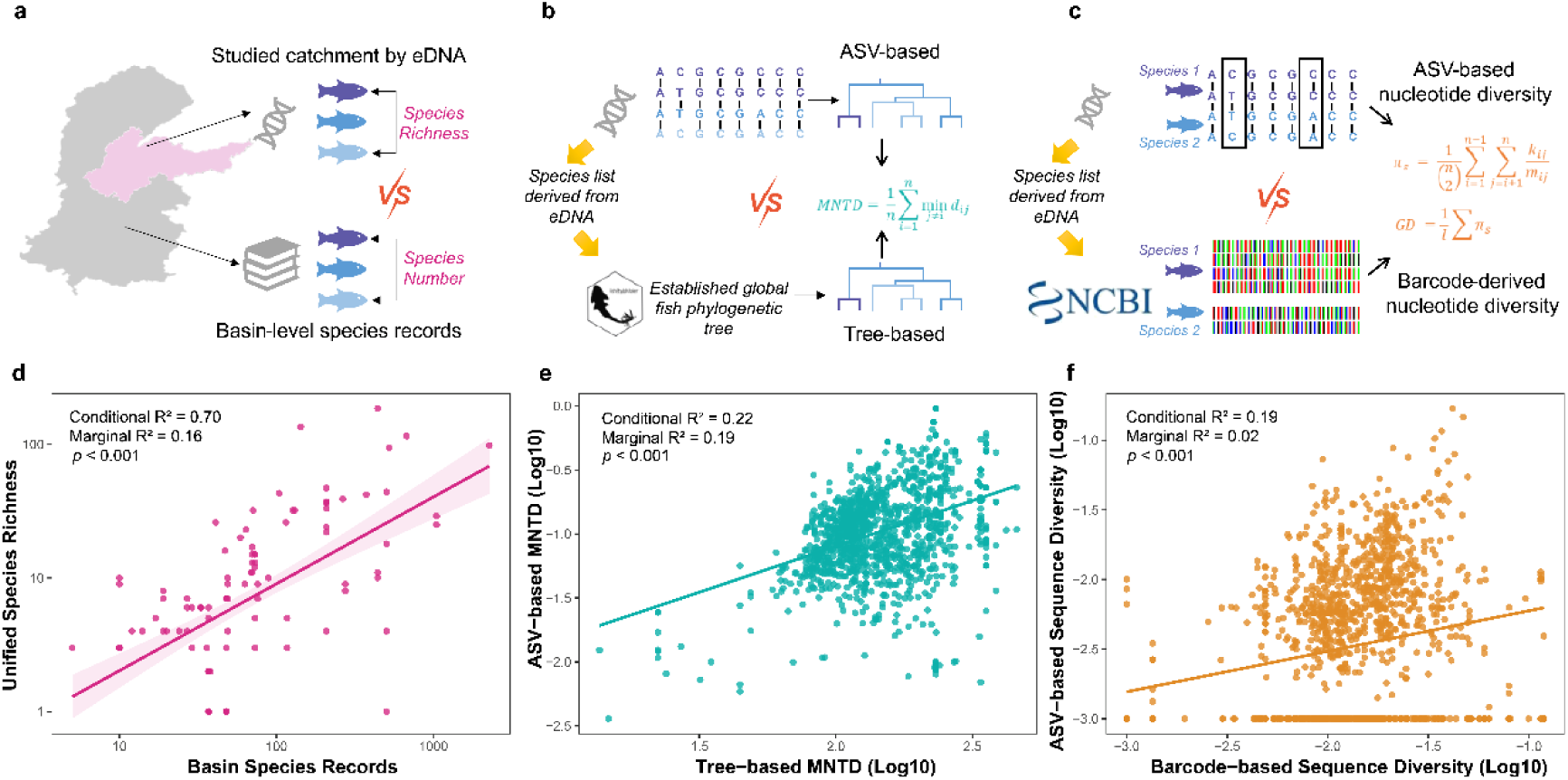
Assessing the ability of eDNA-based data to capture global patterns of multi-faceted fish biodiversity. Conceptual diagrams illustrating the validation framework for three biodiversity facets, including taxonomic (a), phylogenetic (b), and genetic (c) diversity. Along these lines, (d) relationship between eDNA-derived species richness (unified datasets) at the river catchment scale and recorded species numbers from historical data at the corresponding river basin scale are given. Further (e), comparison of ASV-based MNTD with tree-based MNTD derived from a global freshwater fish phylogeny. (f) Comparison of genetic sequence diversity estimated from within-species sequence variation with barcode-based genetic sequence diversity derived from COI sequences in NCBI. All relationships were modelled using generalized linear mixed models. Shaded areas indicate 95% confidence intervals.

### Drivers of multi-faceted biodiversity

Drainage characteristics, encompassing catchment area, climate and human activity intensity, shape biodiversity facets, yielding consistent yet facet-specific patterns (Figure 3 and Supplementary Table 2). Both unified and individually-reported species richness were consistently and positively associated with catchment area (Estimate = 0.14 for unified and 0.39 for individually-reported, *p* < 0.001) and temperature (Estimate = 0.15 *p* = 0.002 for unified; estimate = 0.19, *p* < 0.001 for individually-reported). Spatial autocorrelation had contrasting effects between data types, being positive for unified richness (Estimate = 0.15, *p* = 0.001) but negative for original individually-reported richness (Estimate = -0.11, p < 0.001). The effect of human activities–approximated by intensity of nighttime light–also differed, showing a negative relationship with unified species richness (Estimate = -0.11, *p* = 0.002) but a positive relationship with individually-reported species richness (Estimate = 0.13, *p* < 0.001). Both models revealed a strong negative interaction between catchment area and human activities (Estimate = -0.16 and -0.22, *p* < 0.001), with likelihood ratio tests confirming its significance (Supplementary Table 3, χ^2^ = 30.57 for unified and 71.56 for individually-reported, both *p* < 0.001). Additionally, a positive interaction between catchment area and temperature was significant for eDNA-based richness (Estimate = 0.08, *p* = 0.036; χ^2^ = 8.25, *p* = 0.004), but not for individually-reported richness (χ^2^ = 2.42, *p* = 0.120).

**Figure 3.**
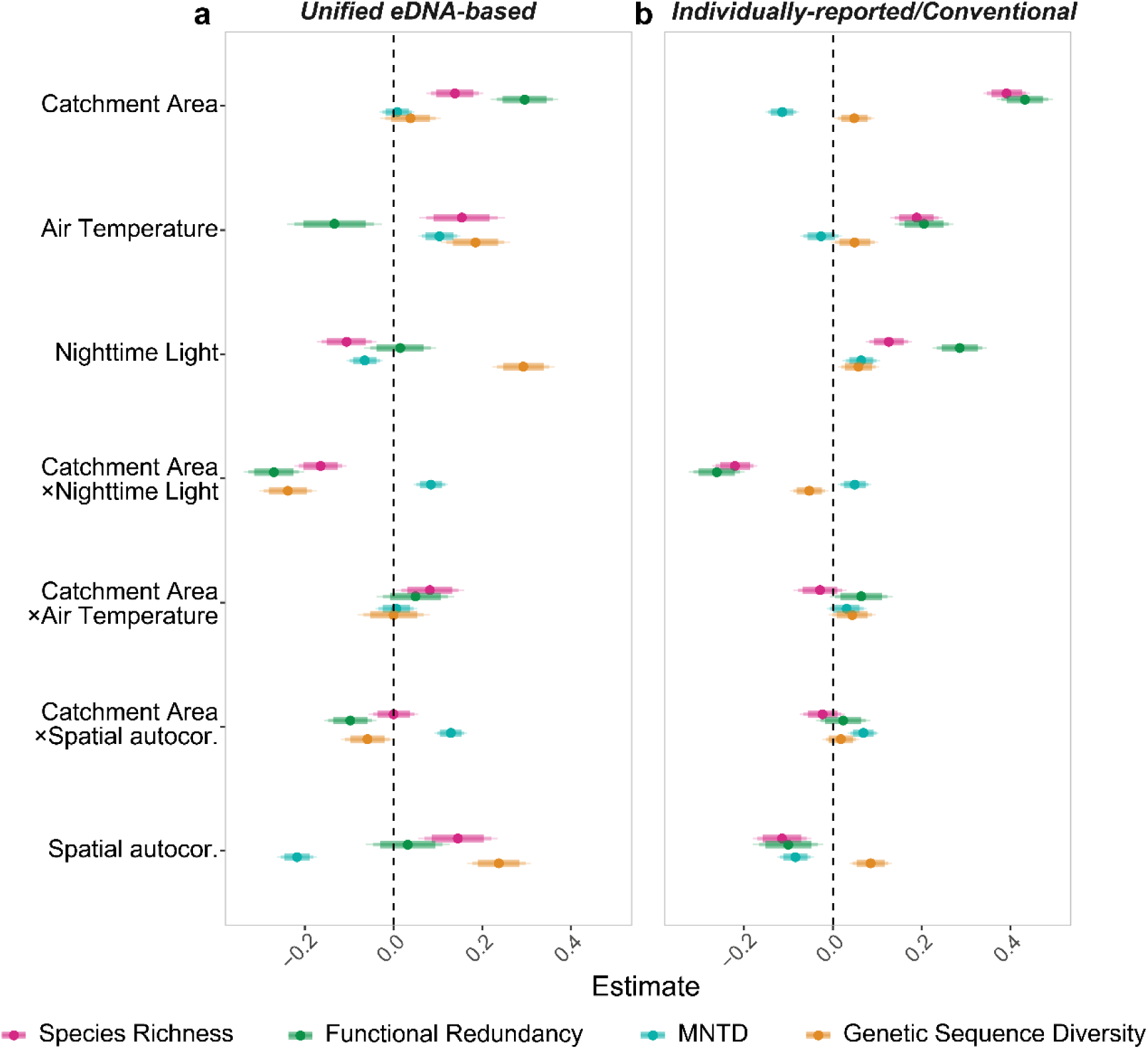
Standardized effect sizes of drainage characteristics on multi-faceted fish biodiversity. Drainage characteristics (including catchment area and its interactions with environmental variables) were modeled across four biodiversity facets (species richness, functional redundancy, MNTD, and genetic sequence diversity). Estimates were obtained using generalized linear mixed models. (a), Models based on all unified eDNA datasets (species richness: 1244 sites; functional redundancy: 1111 sites; MNTD: 1067 sites; genetic sequence diversity: 1199 sites). (b), Models based on the original individually-reported or conventional diversity metrics (species richness: 1601 sites; functional redundancy: 1434 sites; MNTD: 1023 sites; genetic sequence diversity: 1116 sites). Points represent estimated fixed effects, with three bars indicating 80% (thick), 90% (medium), and 95% (thin) confidence intervals. The vertical dashed line indicates zero effect.

Functional redundancy exhibited a broadly similar pattern. It was strongly and positively associated with catchment area (Estimate = 0.30 for unified and 0.43 for individually-reported, *p* < 0.001), while the negative interaction with human activities (Estimate = -0.27 and -0.26, *p* < 0.001) was highly significant according to likelihood ratio tests (χ^2^ = 61.35 for unified and 65.71 for individually-reported, both *p* < 0.001). Temperature effects were again divergent between datasets, being negative for unified data (Estimate = -0.13, *p* = 0.015) but positive in the individually-reported data (Estimate = 0.21, *p* < 0.001). Notably, the main effect of temperature was not significant for unified functional redundancy (χ^2^ = 0.69, *p* = 0.407) but highly significant for individually-reported functional redundancy (χ^2^ = 19.77, *p* < 0.001). A positive interaction between catchment area and temperature was significant for both datasets (χ^2^ = 8.42, *p* = 0.004 for unified; 18.52, *p* < 0.001 for individually-reported), suggesting that the functional benefits of larger catchments were amplified under warmer conditions. Spatial autocorrelation had a significant negative effect only in the individually-reported functional redundancy (Estimate = -0.10, *p* = 0.013; χ^2^ = 27.94, *p* < 0.001) but not in the unified metric (*p* = 0.505).

Phylogenetic diversity, measured by MNTD, was predominantly shaped by spatial processes. ASV-based MNTD decreased significantly with spatial autocorrelation (Estimate = -0.22, *p* < 0.001; χ^2^ = 59.89, *p* < 0.001) and increased with temperature (Estimate = 0.10, *p* < 0.001; χ^2^ = 32.99, *p* < 0.001). Catchment area showed no significant main effect on ASV-based MNTD (Estimate = 0.01, *p* = 0.680; χ^2^ = 8.00, *p* = 0.005), but a strong positive interaction between catchment area and spatial autocorrelation (Estimate = 0.13, *p* < 0.001; χ^2^ = 59.89, *p* < 0.001) indicated that larger catchments mitigated spatial lineage clustering. Tree-based MNTD exhibited similar patterns, with a significant negative effect of spatial autocorrelation (Estimate = -0.08, *p* < 0.001; χ^2^ = 15.67, *p* < 0.001) and a negative main effect of catchment area (Estimate = -0.11, *p* < 0.001; χ^2^ = 27.23, *p* < 0.001). The positive interaction between catchment area and spatial autocorrelation was also significant (Estimate = 0.07, *p* < 0.001; χ^2^ = 15.67, *p* < 0.001). Temperature had no significant effect on tree-based MNTD (*p* = 0.269; χ^2^ = 9.91, *p* = 0.002), despite being statistically significant by likelihood ratio tests, suggesting a weak model-level influence.

Genetic sequence diversity revealed broadly consistent directional responses but different magnitudes between ASV-based and barcode-derived metrics. ASV-based genetic sequence diversity increased with spatial autocorrelation (Estimate = 0.24, *p* < 0.001; χ^2^ = 1.90, *p* = 0.168), temperature (Estimate = 0.18, *p* < 0.001; χ^2^ = 5.41, *p* = 0.020), and human activities (Estimate = 0.29, *p* < 0.001; χ^2^ = 68.01, p < 0.001), while catchment area showed a marginal main effect (Estimate = 0.04, *p* = 0.271; χ^2^ = 3.97, p = 0.046). The negative interaction between catchment area and human activities was strongly significant (Estimate = -0.24, *p* < 0.001; χ^2^ = 51.67, *p* < 0.001), indicating that spatial scaling benefits were reduced under high human activity intensity. For barcode-derived genetic sequence diversity, catchment area (Estimate = 0.05, *p* = 0.037; χ^2^ = 7.20, *p* = 0.007), spatial autocorrelation (Estimate = 0.09, *p* < 0.001; χ^2^ = 4.39, *p* = 0.036), and human activities (Estimate = 0.06, *p* = 0.014; χ^2^ = 6.24, *p* = 0.013) all had significant positive effects. The interaction between catchment area and temperature was significant (χ^2^ = 5.31, *p* = 0.021) for barcode-based genetic sequence diversity but not for ASV-based diversity (χ^2^ = 2.14, *p* = 0.144). The negative interaction between catchment area and human activities was consistently significant across both metrics (χ^2^ = 51.67 for ASV-based, *p* < 0.001; χ^2^ = 5.69 for barcode-based, *p* = 0.017).

### Climate and human activity modulation of biodiversity–area relationships

Across all biodiversity facets, two interaction terms consistently emerged as dominant predictors of biodiversity-area relationships (Figure 3 and Supplementary Table 3). The negative interaction between catchment area and human activities (approximated by nighttime light; χ^2^ ranging from 5.69 to 71.56, *p* < 0.05) was pervasive except for MNTD, indicating reduced biodiversity–area scaling under high human activity intensity. In contrast, the positive interaction between catchment area and temperature (χ^2^ ranging from 5.31 to 18.52, *p* < 0.05) was consistently detected for species richness, functional redundancy, and barcode-derived genetic sequence diversity, reflecting stronger biodiversity–area scaling in warmer regions. These results indicated that biodiversity-area relationships might be jointly modulated by climatic and human activity gradients. Therefore, we presented how biodiversity–area relationships were modulated by temperature and human activities across facets.

The relationship between biodiversity–area slopes and temperature exhibited consistent but facet-specific patterns (Figure 4a–d). For species richness and functional redundancy (both unified and original individually-reported), slopes increased significantly with temperature (*p* < 0.001), indicating that warmer climates promote a higher rate of biodiversity accumulation with spatial expansion. Barcode-derived genetic sequence diversity showed a similar positive relationship (*p* < 0.001). In contrast, ASV-based genetic sequence diversity exhibited a positive but non-significant trend (*p* = 0.517). MNTD displayed a distinct pattern with tree-based MNTD exhibiting a marginally significant positive response to temperature (*p* = 0.016), whereas ASV-based MNTD showed a weak and non-significant relationship (*p* = 0.367) with a positive direction. These temperature-driven modulations of biodiversity–area slopes were robust under the resampling procedure, confirming the robustness of the observed patterns (Supplementary Figure 5).

**Figure 4.**
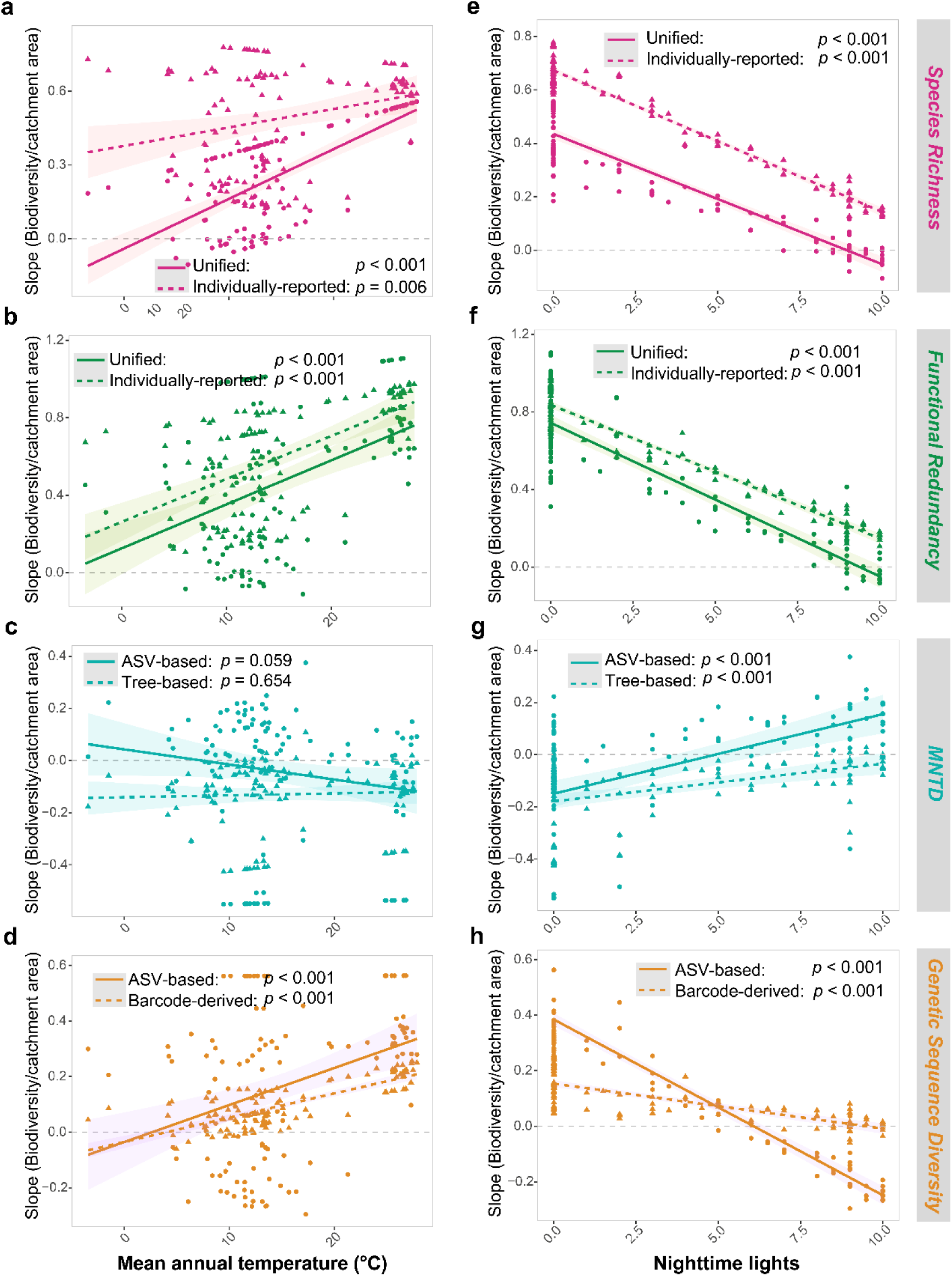
Effects of temperature and human activities on biodiversity–area relationships across river catchments. Relationships between diversity–area slopes estimated from generalized linear mixed models using median mean annual temperature (a–d) and nighttime lights density (e–h) for each river catchments. Each point represents a river catchment. Colored lines are generalized linear model fits for each biodiversity facet. Solid lines represent unified biodiversity metrics, while dashed lines represent original individually-reported or conventional metrics. Shaded areas indicate 95% confidence intervals.

In contrast, the intensity of human activities exerted a broadly negative effect on biodiversity–area slopes across all facets except MNTD (Figure 4e–h). Slopes for species richness and functional redundancy (both unified and individually-reported) declined significantly with increasing human activities (*p* < 0.001), indicating that human activity intensity reduced the rate at which biodiversity accumulates with increasing area. Genetic sequence diversity followed the same pattern, with significant negative relationships for both ASV-based (*p* < 0.001) and barcode-derived metrics (*p* < 0.001). MNTD again exhibited an opposite response, with slopes for both ASV-based and tree-based MNTD increasing significantly with human activities (*p* < 0.001), suggesting that phylogenetic community structure became increasingly homogenized across space in more heavily disturbed catchments. These negative effects of human activities on biodiversity–area slopes were also robust under the downsampling and reshuffling procedures (Supplementary Figure 6).

### Sensitivity to human activities along river network position

The significant interaction between catchment area and human activities not only reflected the modulation of biodiversity–area relationships by human activity intensity but also revealed substantial variation in the responses of fish biodiversity to human activities within river networks.

Biodiversity responses to human activities exhibited clear and facet-specific dependence on river network position (Figure 5). For species richness, functional redundancy, and genetic sequence diversity, the slopes of biodiversity–human activities relationships were significantly more negative in smaller than larger catchments (*p* = 0.010 for barcode-derived genetic sequence diversity, and *p* < 0.001 for others). This indicated that the negative impacts of human activities were stronger in larger catchments, typically representing downstream sections of river networks, whereas headwater streams with smaller catchment areas experienced comparatively weaker negative impacts. In contrast to the other biodiversity facets, the slope of the MNTD– human activities relationship increased significantly with catchment area (*p* < 0.001; tree-based *p* = 0.001), suggesting that the negative effects of human activities on phylogenetic diversity were strongest in smaller catchments, leading to pronounced phylogenetic homogenization, but these effects weakened as catchment area increased, even shifting toward neutral or positive responses in larger catchments. These patterns were consistent and statistically robust, as revealed by the downsampling-based modeling combined with reshuffling-based *p* value estimation (Supplementary Figure 7).

**Figure 5.**
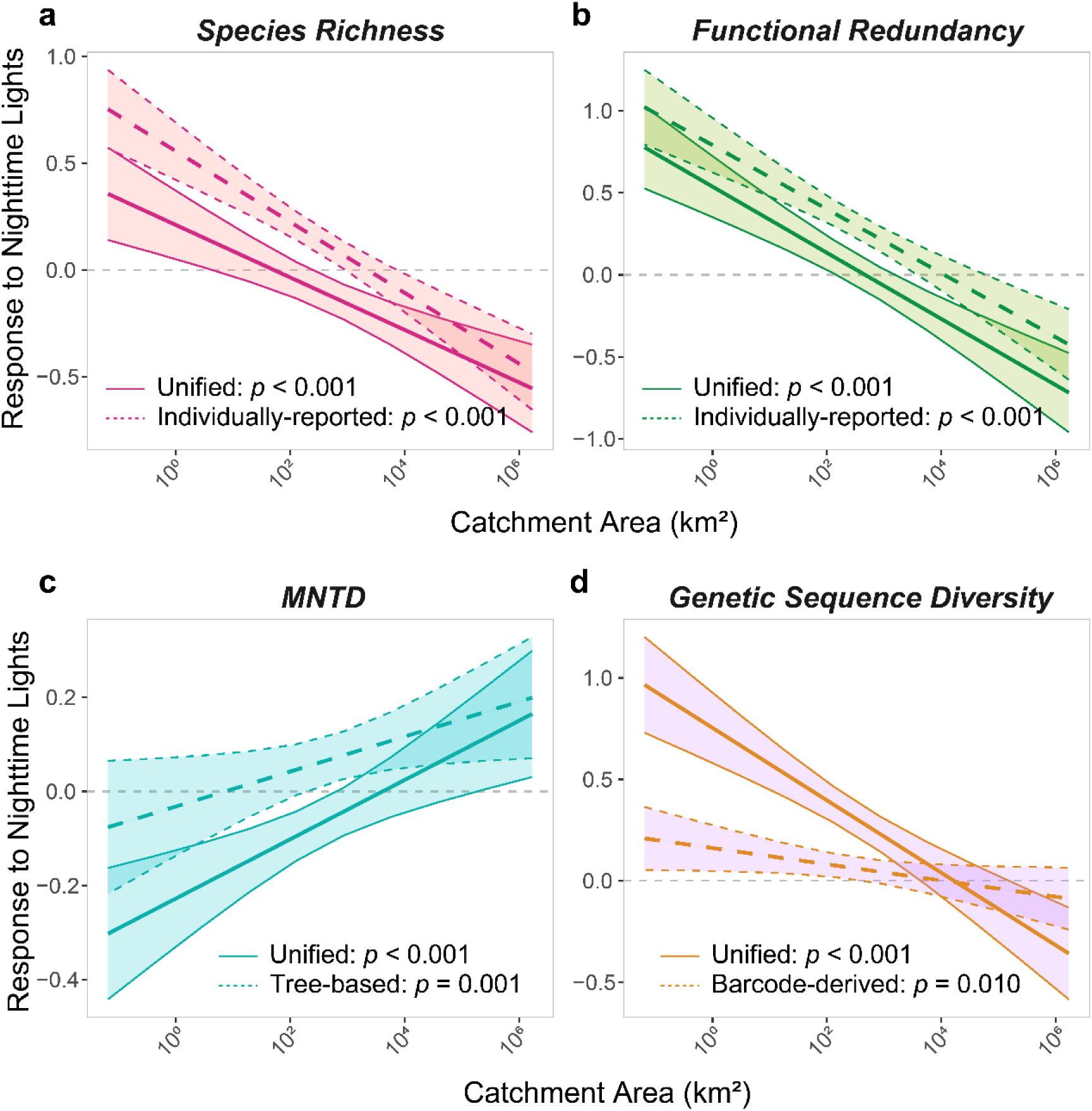
Responses of fish diversity to human activities across river network positions. Relationships between catchment area and diversity–human activities slopes were estimated from generalized linear mixed models based on 1000 repeated 70% downsampling replicates. Human activities were represented by nighttime light data, and catchment area was used as a proxy for the position in the river network. Solid lines represent the mean marginal effect of catchment area derived from unified diversity metrics, while dashed lines correspond to the original individually-reported or conventional metrics. The shaded area indicates the 95% confidence intervals of the marginal effects. Slopes reflect the strength and direction of biodiversity responses to nighttime light intensity. Negative slopes indicate stronger biodiversity loss with increasing human activities. All *p* values reflect the proportion of reshuffled models (1000 times) with estimated effects exceeding the median of the downsampling distribution.

## Discussion

As one of the first global-scale assessments of riverine fish biodiversity using unified eDNA data collected from 1818 sampling sites across 113 river systems, we derived diversity patterns across taxonomic, functional, phylogenetic, and genetic facets of fish across rivers systems globally. We find that eDNA-based biodiversity assessments, based on a meta-analysis of individual studies, capture regional to global riverine fish diversity patterns that can be used for attributing state and change of biodiversity and thus provide guidance for conservation measure. Further, we demonstrate that the common method of eDNA analysis captured associations of biodiversity patterns with environmental drivers (temperature and catchment area) as well as human activity impacts. Biodiversity–area relationships were strongly shaped by drainage characteristics, with an amplification of this scaling along a temperature gradient, while human activities were disrupting it. In parallel, biodiversity responses to human activities exhibited clear spatial heterogeneity. Species richness, functional redundancy, and genetic sequence diversity experienced stronger negative impacts in larger catchments, whereas phylogenetic diversity showed the strongest negative effects in smaller catchments. These findings revealed that spatial structure embedded in river networks fundamentally controls not only biodiversity scaling but also the spatial heterogeneity of biodiversity vulnerability to human activities.

Environmental DNA provides a powerful approach to assess multi-faceted fish biodiversity across global rivers simultaneously. A growing body of research has explored global patterns of freshwater fish biodiversity, most often focusing on taxonomic diversity and its environmental drivers^10,43–46^. Some studies have extended to functional and phylogenetic diversity by incorporating trait data or applying well-established global fish phylogenies^10,44^, advancing our understanding of freshwater biodiversity distribution and threats^47,48^. However, most existing efforts rely on datasets with coarse spatial resolution or biased geographic representativeness, with limited information on genetic diversity^5,49–51^. We demonstrate that eDNA data, when processed through a standardized analytical framework, can overcome many of these constraints by enabling simultaneous assessments of taxonomic, functional, phylogenetic, and genetic sequence diversity, thereby providing a valuable supplement to existing global data on fish biodiversity distribution^29,52^. Despite the short length and locus-specific nature of eDNA barcodes, our analysis shows that they can recover global patterns in species richness, as well as approximate phylogenetic and sequence diversity structures consistent with existing knowledge for fishes. While we do not aim for complete spatial or taxonomic coverage, we provide a proof-of-principle that such harmonized, multi- faceted assessments are feasible and can yield generalizable patterns. As the availability of eDNA data continues to grow, this framework can be readily scaled up, offering a path toward increasingly comprehensive and comparable global biodiversity assessments^41^.

Moreover, our unified analysis of eDNA datasets allows us to attribute fish biodiversity distribution to both natural and human-related drivers, highlighting variations in their effects across different biodiversity facets. The biodiversity–area relationship, a classic pattern in riverine ecology, exhibits contrasting trends across regions and studies, with positive^23^, negative^20^, or non-significant^21^ relationships reported. Our results demonstrate that species–area relationship slopes are strongly influenced by climate change and human activities. In line with the species-energy hypothesis, we find that species–area relationship slopes (except MNTD) are generally steeper in tropical regions than in temperate ones, confirming previous expectations^53^. We further show that human activities generally weakens the species–area relationship, which is consistent with previous studies suggesting that human impacts reduce the likelihood of detecting rare, vulnerable, or habitat-specialist species, thereby flattening species– area relationship slopes^23,54^. Notably, this weakening effect is facet-dependent, with MNTD exhibiting a weaker or even negative relationship with catchment area, reflecting the loss of unique lineages in larger catchments (e.g., in the western Amazon^55^). Such facet-dependent responses highlight the need for multi-faceted biodiversity assessments, as different dimensions of biodiversity exhibit fundamentally distinct and scale-dependent responses to environmental change^33^.

Integrating previously isolated eDNA datasets enables standardized, multi-faceted assessments of fish biodiversity and fosters global collaboration for biodiversity monitoring. Given the massive global human impacts on biodiversity^11^, we need rapid assessment and attribution of state and change of biodiversity at both regional and global scales^38^. Such monitoring and general assessments are also central to many of the goals of the Kunming-Montreal Global Biodiversity Framework ^36^. Yet, achieving such common assessments has hitherto been challenged by methodological constraints, and many datasets—once collected—remain isolated. In this study we show that these common assessments can be achieved for eDNA datasets. By integrating previously isolated datasets through eDNA, we carried out a unified analysis that enabled us to capture multi-faceted distribution patterns of fish biodiversity that are consistent with conventional biodiversity data, despite methodological differences. Specifically, the use of eDNA, not only enabled standardized sampling^41^, but also allowed us to integrate the raw data—millions of sequences generated from environmental samples—into a coherent biodiversity synthesis. Beyond technical standardization, our study unites eDNA researchers globally and demonstrates the potential of unified analyses of isolated eDNA datasets. We therefore encourage the development of more adaptable and sustainable data-sharing and collaboration networks, providing a pathway to contribute meaningfully to global biodiversity monitoring and conservation efforts.

In summary, using an extensive integration and first unified analysis of 58 individual datasets using eDNA globally, we demonstrate that this method provides an effective and scalable approach for capturing riverine fish biodiversity across taxonomic, functional, phylogenetic, and genetic facets, and in particular allows standardized analyses under common frameworks, including identification, taxonomy and sampling efforts. This integrated and unified analysis of eDNA patterns reliably revealed how biodiversity patterns were shaped by major climate factors and human activities, and thus can be directly used to understand and manage freshwater biodiversity. These insights underscored the necessity for conservation frameworks to explicitly account for the influence of climate and human activities on biodiversity patterns and the contrasting vulnerabilities of biodiversity facets.

## Methods

### Fish eDNA data and biodiversity metrics

We compiled a global eDNA fish dataset (for detail see Ref. ^42^), which integrated both original individually-reported and unified fish community data from 113 river catchments worldwide, encompassing 1818 sampling sites (Figure 1a). The data set included both published and unpublished individual eDNA studies that assess riverine fish diversity across six continents, covering the time period 2012–2023. For details on the different contributing datasets, site, geographic and further information, see Supplementary Table 4. For each site, we obtained either (i) an original individually-reported species richness or community matrix extracted from previous studies, (ii) a unified community matrix generated from standardized processing of raw sequencing data, or (iii) both original individually-reported and unified data.

To quantify biodiversity across river catchments, we assessed four complementary facets, including taxonomic, functional, phylogenetic, and genetic (sequence) diversity. Species richness was calculated from both the original individually-reported and the unified datasets based on presence records, with 51 out of 113 river catchments having both values available. Functional diversity was derived for both datasets using species-level morphological traits obtained from the FISHMORPH database^56^. All trait values are continuous and were therefore standardized, and dimensionality reduction was conducted using principal component analysis (PCA) with Euclidean distance^57^. The number of retained axes was determined by parallel analysis implemented in the “paran” R package^58^. Functional redundancy, which quantifies the degree of trait overlap among species within a community, was computed using the Trait Probability Density (TPD) framework^25^. Species-level trait distributions were modeled using multivariate kernel density estimation via the “ks” package^59^, and aggregated into community-level distributions using the “TPD” package^60^.

Phylogenetic and genetic sequence diversity was assessed only for unified datasets, as these were the ones for which raw sequencing data were available, which was required to generate amplicon sequence variants (ASVs). For phylogenetic diversity, ASV sequences were aligned using MUSCLE with default parameters^61^, and phylogenetic trees were constructed from the aligned sequences using the *make_phylogeny.py* script in QIIME. We then calculated mean nearest taxon distance (MNTD), defined as the average phylogenetic distance between each taxon and its closest non-conspecific relative:

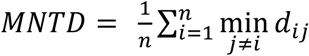

where 𝑑_𝑖𝑗_ denotes the pairwise phylogenetic distance between taxa i and j, and n is the number of taxa in the community. MNTD was calculated using the *mntd*() function from the “picante” package^62^. We also extracted the list of species identified in the unified dataset and retrieved their known phylogenetic relationships from a previously published global phylogeny of freshwater fishes using “FishPhyloMaker” package^63^. Using this species-level phylogeny, we calculated the phylogenetic diversity following the same method as for the tree-based MNTD.

Genetic sequence diversity was quantified as within-species nucleotide diversity (πₛ). This metric primarily captures sequence-level genetic variation, without explicitly considering haplotype richness or frequency distribution^64^. However, given the nature of eDNA metabarcoding data, where accurate genotype resolution is constrained and allele calling is infeasible, πₛ provides an effective, sensitive, and scalable proxy for within-species genetic diversity^35^. In detail, for each species with multiple sequences, we computed the average pairwise sequence divergence as:

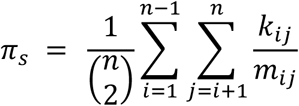

where 𝑘_𝑖𝑗_ is the number of differing base pairs, 𝑚_𝑖𝑗_ is the number of shared base pairs between sequences i and j, and n is the number of sequences per species. Only sequence pairs with more than 50% alignment overlap were retained. The overall genetic sequence diversity (GD) was calculated as the mean nucleotide diversity across all species:

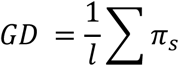

where l is the number of species with valid sequence data. We further obtained COI (cytochrome oxidase I) sequences from NCBI for the species identified in the Unified data, following the approach of Manel et al.^46^. COI was selected for its high resolution in species discrimination. Only deduplicated sequences longer than 600 bp were retained, and πₛ was calculated as described above. We use for each species all available COI data as a first approximation of within-species genetic diversity, acknowledging this does not control for different sampling efforts between species and may not completely reflect a species’ local genetic diversity, but is rather an integrated metric. Given that most ASVs targeted 12S and 16S barcode regions, we also calculated nucleotide diversity based on sequences from these markers to enable cross-marker comparison.

In summary, a total of 661 fish species were included in the unified eDNA datasets for quantifying species richness and functional redundancy (Supplementary Figure 8a). All species were used to calculate tree-based phylogenetic diversity, while genetic sequence diversity was computed for 520 species. The original individually-reported diversity data encompassed 1521 species, although a substantial proportion (517 species) were identified only to the genus level (Supplementary Figure 8b). As a result, functional diversity calculations based on the original individually-reported datasets were limited to 782 species with validated species-level trait information.

### Validation of biodiversity metrics derived from eDNA

We used generalized linear mixed models (GLMMs) to assess whether biodiversity metrics derived from eDNA reflect global patterns of freshwater fish diversity^65^. For taxonomic, phylogenetic, and genetic sequence diversity, we separately modeled the eDNA-inferred values as a function of the corresponding reference values, while accounting for sampling and technical variation (Figure 2a). For taxonomic diversity, catchment-level species richness estimated from eDNA was modeled as a function of observed richness compiled from historical occurrence records ^48^. For phylogenetic and genetic sequence diversity, MNTD and nucleotide diversity derived from ASVs were compared respectively with MNTD calculated from a species-level phylogeny and with COI-based genetic sequence diversity obtained from public databases. Filtered water volume was included as a fixed covariate in all models to account for differences in sampling effort. Amplification-related variations were modeled by including either primer identity or barcode region as a random effect. All response and predictor variables were log-transformed (base 10) prior to model fitting. Models were fitted using the “glmmTMB” package in R^66^.

### Delineation of river catchments and extraction of drainage characteristics

To better capture the river catchment characteristics, we employed the widely used D8 algorithm and the Hydrography90m flow direction map (three arc second resolution, ∼90 m at the equator) to calculate the upstream catchment for eDNA sampling sites^67^. In brief, the value of a pixel in the flow direction map indicates the water flow direction to eight adjacent pixels. To begin with, river channels were directly extracted from the flow accumulation map of Hydrography90m, with which we manually checked and calibrated the coordinates of all the eDNA sampling sites. By tracking along the water flow direction map, one can determine whether a terrestrial pixel ultimately flows through a sampling site and then calculate its water flow distance. These terrestrial pixels that flowed through the sampling site were then connected to produce a site catchment map (Supplementary Figure 9). Because of worldwide spatial coverage of sampled rivers, we used the haversine formula to estimate the length and width of map pixels for better accuracy in the flow distance computation. In addition, we calculated the catchment area for all the sampling sites based on these catchment maps.

We used the WorldClim version 2.1 climatic data (30-second resolution) for 1970— 2000 to assess the climatic effects on global fish distribution patterns (website: https://www.worldclim.org/data/worldclim21.html)^68^. Among all the bioclimatic variables, we adopted mean annual temperature to represent the energy of the environment and extracted these values from raster layers at the sampling sites. Nighttime light data was used as a typical proxy for human activity intensity. We directly used the nighttime light layer from Venter *et al.*, 2016^69,70^ developed based on the Defense Meteorological Satellite Program Operational Linescan System (DMSP-OLS) nighttime light data. To better represent the human activities from the surrounding upstream area, we extracted the nighttime light raster values within a 4 km upstream flow distance buffer for each sampling site and the calculated the median as the proxy for human activities.

### Statistical modelling of biodiversity responses

To evaluate how drainage characteristics influence fish biodiversity across river networks, we used generalized linear mixed models (GLMMs) with a beta distribution and logit link function, implemented via the “glmmTMB” package in R^66^. The response variables included species richness, functional redundancy, MNTD, and genetic sequence diversity. These biodiversity metrics exhibited right-skewed distributions and varied in scale (Figure 1 and Supplementary Table 5). Although their measurement units and ranges vary, all metrics share a common statistical property in that they are continuous and constrained within a fixed interval. This supports the use of beta distribution as a consistent and appropriate modelling approach across biodiversity metrics. Accordingly, prior to modelling, all response variables were log-transformed to stabilize variance and subsequently rescaled to the interval between zero and one. The beta distribution was selected for its capacity to accommodate continuous, bounded, and asymmetric data structures, offering improved flexibility and robustness compared to Gaussian alternatives^71^. Logit link allowed covariate effects to be estimated on an unbounded latent scale while preserving the ecological interpretability of predictions within a bounded outcome space^72^.

The null model accounted for amplification and sampling variation by including sampling volume as a covariate, spatial autocorrelation as a fixed effect, and a random intercept for either primer or barcode region. Spatial autocorrelation was defined as the average minimum distance between sampling sites within each river catchment, reflecting local clustering while accounting for heterogeneity in catchment extent. The basic model added catchment area as a proxy for network position, assuming biodiversity scales with drainage area due to cumulative dispersal, environmental filtering, and habitat complexity. An interaction between catchment area and spatial autocorrelation was included to account for differences in the spatial structure of sampling within catchments. The full model further incorporated mean annual temperature and nighttime light intensity as proxies for climate and human activity intensities, respectively. Interactions with catchment area enabled evaluation of whether spatial scaling relationships vary along climatic or human activity gradients.

Models were fitted independently for each biodiversity metric. All predictor variables were standardized by mean-centering and scaling to unit variance, with catchment area log10-transformed prior to standardization (Supplementary Figure 10). Model fit was evaluated by plotting fitted versus observed values for each diversity metric (Supplementary Figure 11). Models captured the overall variation reasonably well (GAM: R^2^ = 0.15–0.43), consistent with typical expectations for noisy ecological data. Model performance was further assessed using AIC, BIC, log-likelihood, and both marginal and conditional R² values. Nested models were compared using likelihood ratio tests to evaluate whether the inclusion of additional terms significantly improved model fit (Supplementary Table 6)^73^. Based on these comparisons, primer identity was used as the random intercept for species richness and functional redundancy, while barcode region was used for phylogenetic and genetic sequence diversity (Supplementary Figure 12). Fixed-effect estimates were extracted to assess the relative importance of predictors across biodiversity metrics.

### Prediction and interpretation of model outputs

To examine the climatic and human activity dependence of biodiversity scaling with catchment area, we performed a series of marginal slope predictions based on the full model structure. First, for each biodiversity metric, we evaluated how climatic variable (mean local annual temperature) and intensity of human activities (approximated by nighttime light intensity) modulate the relationship between biodiversity and catchment area. Median values of annual temperature, nighttime light intensity and spatial autocorrelation were calculated for each river catchment by aggregating all sampling sites within the corresponding drainage area. These aggregated values defined the climatic and human activity conditions under which the biodiversity–area relationship was evaluated. Using the *emtrends()* function from the “emmeans” package^74^, marginal trends in biodiversity as a function of catchment area were predicted for each catchment.

To evaluate the uncertainty of slope estimates, we repeatedly performed full model fitting followed by marginal slope estimation, each based on a random subsample comprising 70% of the total sites. This process was repeated 1,000 times. In a parallel analysis, we assessed whether responses of biodiversity to human activities vary along the river network by predicting the marginal slope of biodiversity in response to nighttime light intensity across a standardized gradient of catchment area. A grid of values spanning the observed range of catchment size was used to generate these predictions.

To test whether observed biodiversity–area scaling arises from genuine ecological patterns rather than from spatial autocorrelation in covariates, we implemented a covariate reshuffling procedure repeated 1,000 times. In each iteration, temperature and nighttime light intensity were randomly reassigned among sampling sites. Slope estimates under reshuffling were compared to those from downsampling-based predictions, and *p* values were calculated as the proportion of reshuffles with slopes exceeding the median from the downsampling procedure (Supplementary Figure 13)^75^.

## Acknowledgments

This study is supported by the National Key Research and Development Program of China (2022YFC32021001, 2021YFC3201003) and National Natural Science Foundation of China (42507382). We thank all the many people involved in the field and laboratory work of the respective studies we base our reanalysis on. We also thank all researchers who provided comments and information on the published dataset. Funding is from the Swiss National Science Foundation, grants 310030_197410 and 31003A_173074 (to FA). The views expressed in this article are those of the author(s) and do not necessarily represent the views or policies of the U.S. Environmental Protection Agency.

## Author Contributions

Y.Z. and F.A. conceived and led the study. Y.Z. collected and analyzed the datasets, and wrote a first version of the paper, under the supervision of F.A. and X.Z. H.Z. helped with delineation of river catchments and extraction of drainage characteristics. All other authors contributed data. All authors commented on drafts of the paper.

